# In Vitro Culture Alters Cell Lineage Composition and Cellular Metabolism of Bovine Blastocyst

**DOI:** 10.1101/2023.06.09.544379

**Authors:** Hao Ming, Mingxiang Zhang, Sandeep Rajput, Deirdre Logsdon, Linkai Zhu, William B Schoolcraft, Rebecca Krisher, Zongliang Jiang, Ye Yuan

## Abstract

Profiling transcriptome at single cell level of bovine blastocysts derived in vivo (IVV), in vitro from conventional culture medium (IVC), and reduced nutrient culture medium (IVR) has enabled us to reveal cell lineage segregation, during which forming inner cell mass (ICM), trophectoderm (TE), and an undefined population of transitional cells. Only IVV embryos had well-defined ICM, indicating in vitro culture may delay the first cell fate commitment to ICM. Differences between IVV, IVC and IVR embryos were mainly contributed by ICM and transitional cells. Pathway analysis by using the differentially expressed genes of these non-TE cells between groups pointed to highly active metabolic and biosynthetic processes, with reduced cellular signaling and membrane transport in IVC embryos, which may lead to reduced developmental potential. IVR embryos had lower activities in metabolic and biosynthetic processes, but increased cellular signaling and membrane transport, suggesting these cellular mechanisms may contribute to the improved blastocyst development compared to IVC embryos. However, the IVR embryos had compromised development when compared to IVV embryos with notably over-active membrane transport activities that led to impaired ion homeostasis.

**Summary Statement:** Single-cell transcriptomic analysis of bovine blastocysts produced in vivo, and in vitro in conventional and reduced nutrient conditions reveals the effect of culture environments on embryo developmental potential.

## Introduction

In vitro fertilization (IVF) and embryo in vitro production (IVP) technologies have been widely used to treat human infertility and improve reproduction efficiency of agricultural economic species, such as cattle. The number of bovine IVP embryos transferred to a recipient has steadily increased over the years globally. However, the competence of bovine IVP embryos to establish pregnancy is much lower than the embryos produced in vivo, and the optimization of in vitro condition is warranted to improve the embryo competence from IVP. One major obstacle that hampers the further optimization of the bovine embryo in vitro culture condition is the uncertainty of the precise molecular mechanisms regulating bovine early embryonic development and lineage differentiation. Transition from morula to blastocyst involves two cell fate decision events, the first defines inner cell mass (ICM) from the outer trophectoderm (TE) layer and the second results in ICM further segregating into epiblasts (EPI) and primitive endoderm (PE). Using single cell RNA sequencing (scRNA-seq), the distinct molecular profiles of three lineages have been well defined in mice and human (Fan et al., 2020; Liu et al., 2022; Mole et al., 2021; Yan et al., 2013). However, this information is very limited in bovine, where most of current knowledge was obtained using less informative candidate quantitative PCR approaches adopted from other species (Negron-Perez et al., 2017; Wei et al., 2017). Thus, a high throughput sequencing of bovine blastocysts at single cell resolution would inform cell lineage identifies and their molecular signatures of bovine blastocyst.

The optimization of the in vitro culture media is a continued effort to improve the overall efficiency of embryo IVP. Metabolomics analysis revealed that blastocysts from various species consumed significant amount of amino acids from the culture medium, but glucose and other metabolites take-ups were minimal. Significant differences in the metabolic profiles of in vivo-compared with in vitro- produced embryos were also found at the blastocyst stage (Krisher et al., 2015). These findings suggested that the amounts of nutrients present in current culture media far exceed the amounts needed by the embryos. As described in other model systems (Qiu and Schlegel, 2018; Um et al., 2006), such a nutrient overload condition may impair the metabolic homeostasis of the embryos and potentially damage their developmental potential. Recent studies have shown the possibility of producing embryos in vitro using reduced nutrient media in different species, and such condition may result in improved embryo development (Ermisch et al., 2020; Herrick et al., 2020; Santos et al., 2021). However, the molecular mechanism behind such an improvement is still waiting to be explored.

Here, we conducted scRNA-seq analysis of bovine blastocysts derived from conventional in vitro culture condition (IVC) and reduced nutrient condition containing 6.25% of standard nutrient concentrations (IVR), and compared them to their in vivo counterparts (IVV) to address the impact of culture conditions on the transcriptomic variation in bovine blastocysts. We aim to answer the following important questions. First, how does lineage segregation happen at the blastocyst stage, and what are the transcriptomic signatures of each cell lineage identified in a bovine blastocyst? Second, what are the transcriptomic differences between the less competent IVC and IVV embryos, and what are the mechanisms that can explain the beneficial role of IVR condition in improving bovine blastocyst formation?

## Results

### Definition of cell lineages in a bovine blastocyst

Bovine blastocysts were collected at day 8 post fertilization followed by mural trophectoderm (TE) cells removal via microdissection to enrich the cells from the ICM. The remaining ICM cells along with polar TE were enzymatically dissociated into single cells and collected individually for scRNA-seq (Fig. 1A). A total of 195 cells from 19 blastocysts (Table S1) were collected and used for library preparation, obtaining approximately 90.04% of mapping rate in average (Table S2).

**Fig. 1.**
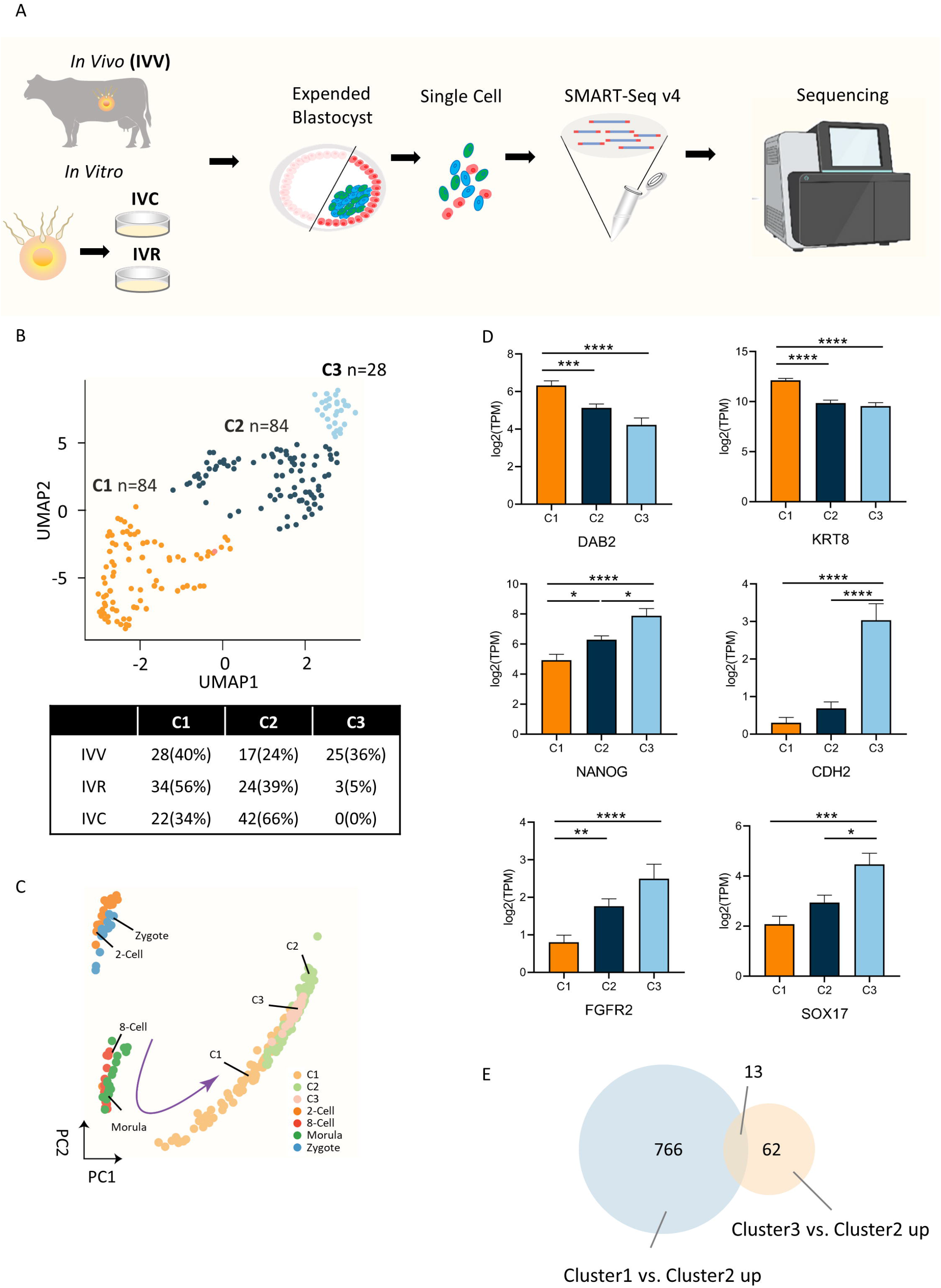
Single-cell transcriptome profiles of *in vivo* and *in vitro* derived bovine blastocysts. A. Scheme of transcriptome profiling of single cells from bovine blastocyst cultured in vivo and in vitro conditions. B. Two-dimensional UMAP representation of cell clusters analyzed using all expressed genes, as well as the cell number/ percentage in each cluster and in each. C. Principal component analysis (PCA) of pseudo bulk conversion of blastocyst cells from different lineages in this project and previous data of embryos from zygote to morula stage (Zhao et al., 2022). D. Log2 TPM values of defined lineage marker genes in three clusters (top lane as trophectoderm markers, middle lane as EPI markers, bottom lane as PE markers). E. Venn diagram of highly expressed genes in cluster 1 and cluster 3 when compared with cluster 2, respectively. All data are presented as the mean ± SEM. * P < 0.05, ** P < 0.01, *** P < 0.001, **** P < 0.0001 from one-way ANOVA followed by Tukey’s multiple comparisons test.

Following transcriptome assembly and quantification of gene expression, we performed two-dimensional principal component analysis (PCA) (Fig. S1A), t-distributed stochastic neighbor embedding (t-SNE) (Fig. S1B), and uniform manifold approximation for dimension reduction (UMAP) using all genes expressed to examine the clustering of all cells (Fig. 1B). As suggested by earlier studies, we expected to visualize the separation of three lineages, including EPI, PE, and TE in our analysis (Negron-Perez et al., 2017; Wei et al., 2017). UMAP provided relatively better separation than PCA and t-SNE (Fig. 1B). Additionally, when single cell RNA-seq data from earlier stage embryos were included, we found that cells from blastocyst and earlier stages were orderly arranged on the PCA plot, suggesting a sequential developmental trajectory from zygote until lineage specification during blastocyst stage (Fig. 1C).

By examining the lineage marker gene expression in each cell cluster (C1, C2, and C3), C1 was found to have more abundant expression of signature genes for TE (*DAB2, KRT8, GATA2, SFN*) compared to C2 and C3, suggesting that C1 cells represented TE lineage (Fig. 1D top panel, Fig. S1D). The signature genes for EPI, such as *NANOG*, *FGF4*, *CDH2*, *SAV1*, were more abundant in C3 cells, and intriguingly, many PE markers, such as *SOX17, PDGFRA, FGFR2, ALPL,* were also enriched in C3 cells, suggesting that cells in C3 cluster were committed to become ICM (Fig. 1D middle and bottom panels, Fig. S1E, F). This result also suggested that the segregation of EPI and PE has not readily accomplished in day 8 bovine blastocysts yet, which is consistent with the previous finding that the well-defined PE lineage was not established until day 10 post fertilization (Maddox-Hyttel et al., 2003). C2 cluster was a peculiar cell population, as it has significantly higher levels of EPI and PE markers (*NANOG*, *SAV1*, *FGF2*, etc.) when compared with C1 cells (TE) while also has more abundant expression of TE markers (*GATA2*, *PECAM1*, etc.) when compared with C3 cells. The mixed transcriptomic signature of C2 cells suggested that these cells may be in a transitional state that their decision in lineage segregation between ICM and TE was delayed (Fig. 1B, Fig. S1D-F). Surprisingly, when we continuously explored the differentially expressed genes (DEGs) between C1 and C2, or C1 and C3, we found 779 genes were highly expressed in C1 when compared to C2, while only 75 genes were more enriched in C3 when compared to C2, indicating although in a transitional status, C2 was much more similar to C3 (Fig. 1E).

Therefore, C1 cells belong to TE, C3 cells belong to ICM, and C2 cells represent transitional cells prone to become ICM and yet to commit to either ICM or TE (Fig. S1C). When examining the cell cluster distribution for each treatment group (IVV, IVC, and IVR), C3 cells were almost exclusively derived from IVV embryos, suggesting that the in vitro culture condition may delay the first cell fate commitment to ICM (Fig. 1B). Cells from IVV embryos had the least percentage (24%) residing in the transitional state (C2), whereas the majority of cells from IVC embryos were transitional cells (C2, 66%) (Fig. 1B). There was a small percentage of cells derived from IVR embryos committed to ICM (C3, 5%), and the majority of the cells from IVR embryos were committed to TE (56%) (Fig. 1B).

### Novel markers and signaling pathways to identify bovine ICM and TE cell lineages

Current knowledge of the lineage segregation and lineage markers in bovine blastocysts are mainly based on mouse and human studies. However, the lineage markers of bovine ICM and TE may differ from other species but remain elusive. Our dataset provides the opportunity to fill up this knowledge gap. We only chose samples from IVV embryos in this analysis in order to eliminate the bias that may be introduced by in vitro turbulence. When comparing the IVV samples in C1 cluster (TE) with C3 cluster (ICM) (Fig. 2A), 581 TE enriched genes and 560 ICM enriched genes were identified, respectively. These differentially expressed genes include not only well-known lineages markers reported in other species, but also the genes that have not been reported to be associated with lineage specification in other species and may be severed as novel lineage markers for bovine ICM and TE. We first examined the putative lineages markers that were reported in earlier studies by screening a predetermined set of genes using single-cell quantitative PCR (Negron-Perez et al., 2017; Wei et al., 2017). A total of 13 EPI markers, 13 PE markers, and 15 TE markers were selected and their expression between TE and ICM from IVV samples were compared (Fig. 2B). We confirmed that EPI markers (Fig. 2B top panel), including *CDH2, POU5F1, FN1, HNF4A, SAV1, FGF4, NANOG*, *TDGF1*, and PE markers (Fig. 2B middle panel), including *ALPL, FGFR2, HDAC8, HNF4A, ID2, PDGFRA, SOX17, GJA1*, and *RUNX1*, had significantly higher expression levels (p<0.05) in ICM compared to TE. As for TE markers (Fig. 2B bottom panel), *ACTA2*, *SFN, DAB2, KRT8, GATA2, PECAM1* and *GATA3* were more abundant (P<0.05) in TE than in ICM in our dataset. However, the other putative markers were not significantly different between the two cell types, questioning whether these genes can be used as lineage markers for bovine TE and ICM. One explanation for such discrepancy is that only in vitro produced bovine blastocysts were used to identify these putative markers in the earlier studies (Negron-Perez et al., 2017; Wei et al., 2017). Therefore, we examined these invalidated markers in the in vitro samples in the dataset and found that in the IVC condition, *GSC, MBNL3*, *H2AFZ, KDM2B, PRDM14* and *MBNL3* were more abundant in their representing lineages (Fig. S2A). The invalidated lineage markers we found here could be either the outcome of adverse impact from in vitro culture, or the limited sample size in those earlier studies. Furthermore, by using more strict cutoff values (P < 0.01, Fold change >2), 94 genes were obtained as the candidates of lineage marker genes (Table S3) and some of them were shown here (Fig. 2C), this gene set could be served as potential markers in bovine specifically for future studies.

**Fig. 2.**
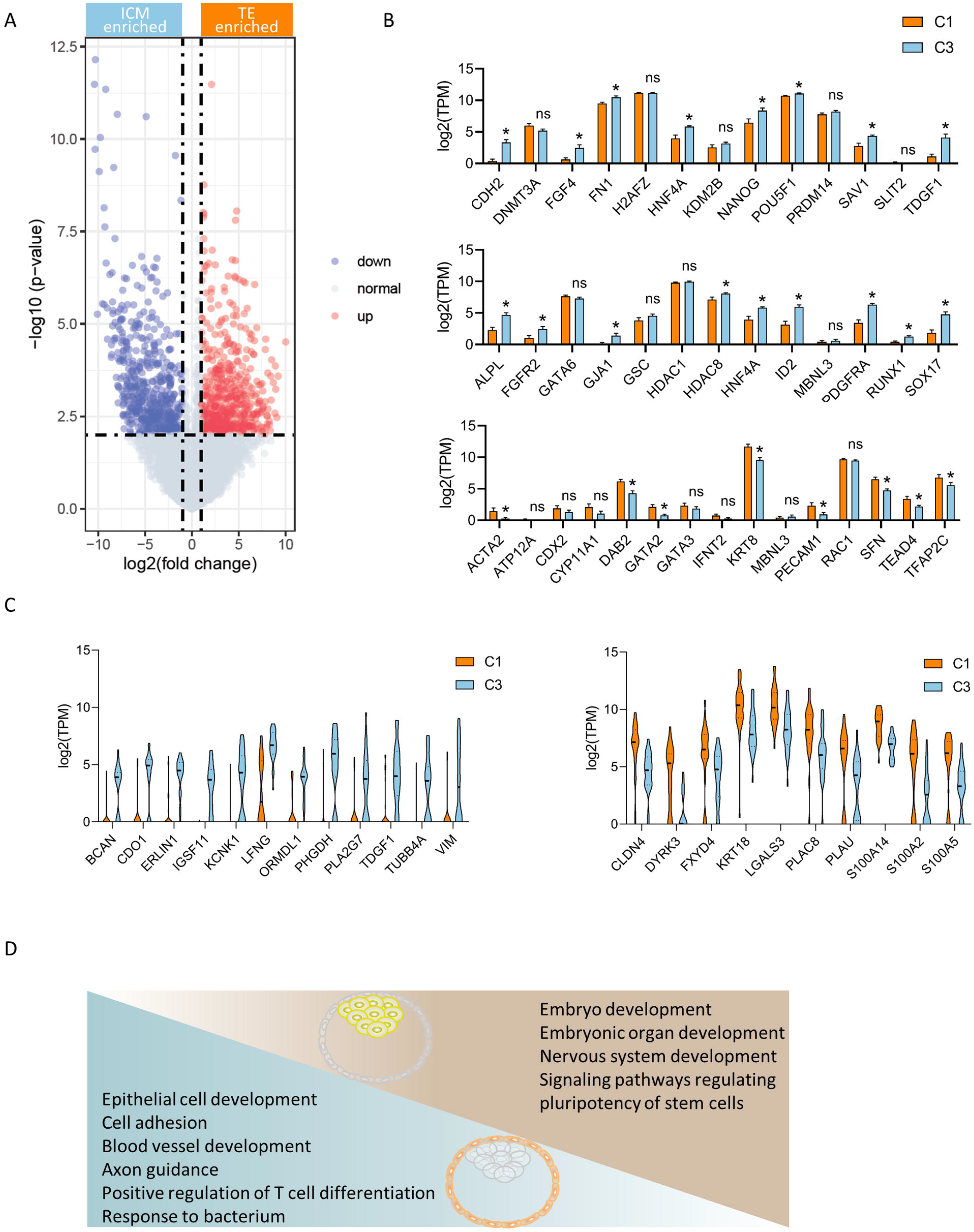
Lineage specification and potential marker genes in bovine blastocyst. A. Volcano plot showing differentially expressed genes (DEGs) between TE cells and ICM cells in IVV embryos. Differentially expressed genes were defined as P < 0.05, Fold change > 2. B. Log2 TPM values of representative potential EPI lineage marker genes (top), PE lineage marker genes (middle), as well as TE marker genes (bottom). C. Log2 TPM values of representative potential ICM (left panel) and TE (right panel) lineage marker genes.D. Top represented GO and KEGG terms for the DEGs between cluster C1 and C3 in IVV embryos. All data are presented as the mean ± SEM. * P < 0.05 from unpaired t-test.

Based on the differentially expressed genes between ICM and TE, we then performed gene ontology (GO) and pathway analysis to identify biological functions and signalings strongly associated with each cell lineage. Again, to remove the potential bias caused by in vitro culture condition, we only involved IVV derived samples for analysis. It was shown that the top GO terms for TE were related to epithelial cell development, cell adhesion, blood vessel development, etc., consistent with its role found in other species. For ICM, the top terms were embryo development, nervous system development, signaling pathways regulating pluripotency, which again consistent with known feature of ICM found in other species (Fig. 2D, Fig. S2B; Table S4). KEGG analysis of these two cell types revealed a distinct metabolic preference of each lineage, in which TE had genes more enriched in pathways related to hippo signaling, calcium signaling pathway, cAMP signaling, Axon guidance, MAPK signaling, oxytocin signaling, whereas ICM is in favor of gap junction, TGF-beta signaling, Wnt signaling, purine metabolism, fatty acid elongation, glycolysis/gluconeogenesis, and fructose and mannose metabolism (Fig. S2C; Table S5).

### The difference in quality between IVV, IVC, and IVR embryos is mainly contributed by non-TE cells

Embryos from all three groups contained a defined TE cell population (C1 cells, in Fig. 1B). However, IVV embryos had a mixture of defined ICM (C3 cells) and transitional cells (C2 cells), whereas IVC and IVR embryos mainly contain transitional cells. As transitional cells are more similar to ICM, we combined these two groups of cells as the non-TE population and compared the transcriptomes of TE population (C1) and non-TE population (C2 and C3) to capture the impact of in vitro culture on development of both lineages.

The IVR condition resulted in significant improvement of blastocyst formation in our previous study (https://www.publish.csiro.au/rd/RDv30n1Ab73). We also validated the benefit of the IVR condition in improving blastocyst formation in this study (Figure S3A), showing much higher blastocyst rate when comparing to IVC. To explore the mechanisms behind such improvement, we first performed Gene Set Enrichment Analysis (GSEA) to examine the developmental potential of TE and non-TE cells, respectively. The ability of TE contributing to embryo and the following placenta development was comparable between cells from IVC and IVV embryos (Fig. 3A). However, the potential of non-TE in embryonic development and implantation process is much higher in IVV embryos than IVC embryos (Fig. 3B). When comparing IVR and IVV embryos, TE lineage showed similar developmental potential between these two groups (Fig. 3C). However, slight difference between IVR and IVV embryos still exists in non-TE lineage, while this gap narrowed significantly compared to the difference between IVC and IVV (Fig. 3D). We also compared IVR and IVC embryos. Similarly, there is no difference in TE lineage development between IVR and IVC embryos (Figure 3E), and again, the developmental potential of non-TE lineage is significantly higher in IVR derived blastocyst than that in IVC (Fig. 3F). These results suggested that the difference in embryo quality between IVV, IVC, and IVR blastocysts is mainly defined by non-TE cells. In addition, IVR embryos may be more competent than IVC embryos and developmentally more similar to IVV embryos.

**Fig. 3.**
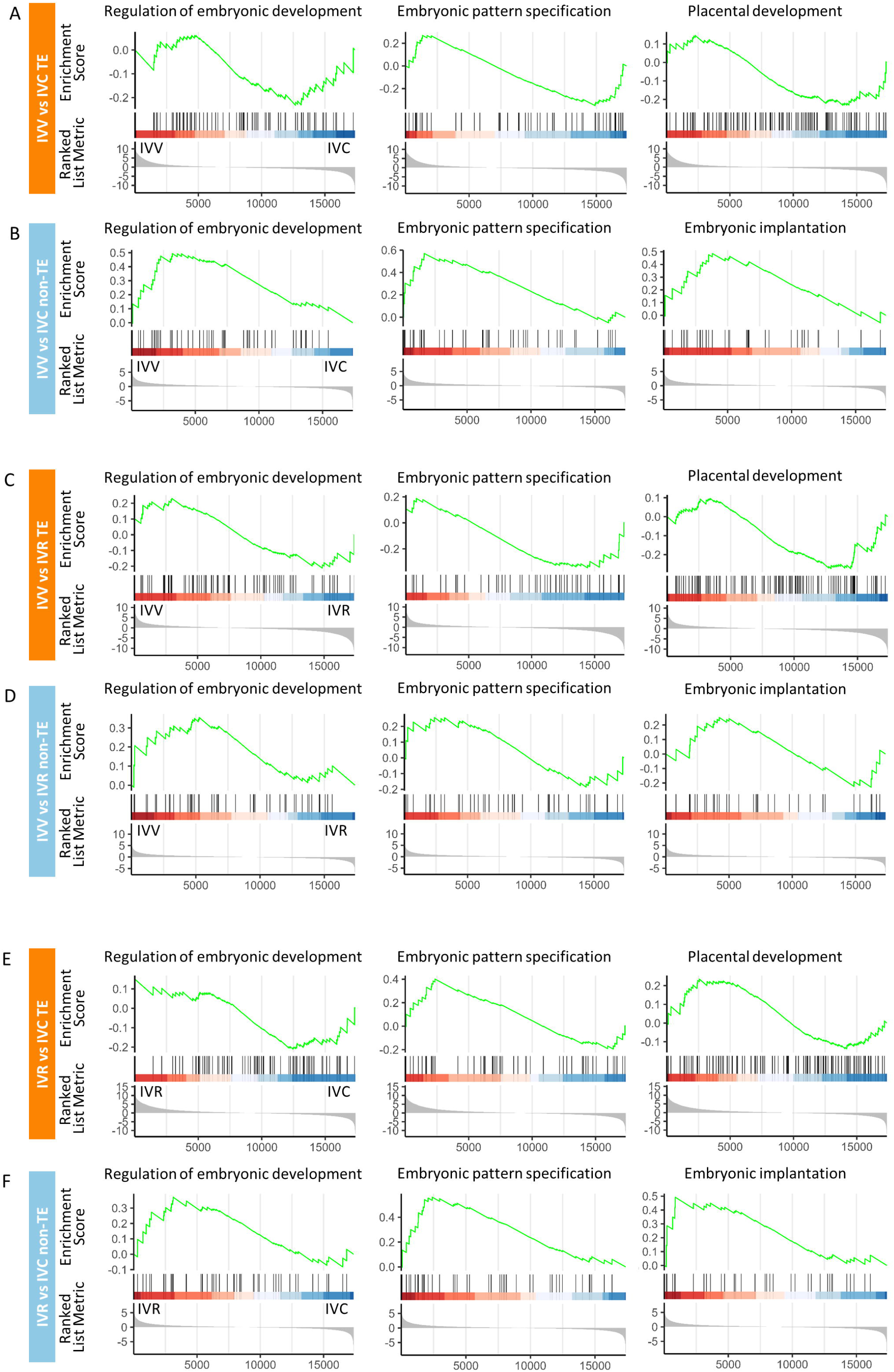
GSEA reveals that difference in embryonic quality is defined by non-TE lineage. A. Gene Set Enrichment Analysis (GSEA) of the enrichment of represented gene ontology (GO) terms regulating embryonic and placenta development in IVV and IVC TE lineage. B. GSEA of the enrichment of represented GO terms regulating embryonic and placenta development in IVV and IVC non-TE lineage. C. GESA of the enrichment of represented GO terms regulating embryonic and placenta development in IVV and IVR TE lineage. D. GESA of enrichment of represented GO terms regulating embryonic and placenta development in IVV and IVR non-TE lineage. E. GESA of enrichment of represented GO terms regulating embryonic and placenta development in IVR and IVC TE lineage. F. GESA of enrichment of represented GO terms regulating embryonic and placenta development in IVR and IVC non-TE lineage.

### Gene ontology analysis of non-TE cells reveals impact of in vitro culture on embryo quality

Based on the fact that non-TE lineage but not TE lineage leads to the difference in quality between IVV, IVC, and IVR embryos, we next unveiled detail information in non-TE lineage specifically. When comparing IVC and IVV embryos, 787 upregulated genes and 489 downregulated genes were found in the TE lineage. For non-TE, 349 genes were upregulated, and 529 genes were downregulated in the IVC compared to the IVV embryos (Fig. S3B). We performed GO analysis on non-TE cells and intriguingly, almost all the GO terms enriched (P<0.05) in either IVC or IVV embryos were able to be summarized into six categories: development and morphogenesis (Development), transmembrane transport (Transport), cellular response and signaling (Cellular Response), biosynthetic process (Biosynthetic), metabolic process (Metabolic), and immune response (Immune) (Fig. 4A). GO terms enriched in IVC embryos were related to biosynthetic and metabolic processes, including amino acid, small molecule, carboxylic acid, lipid biosynthesis, lipid metabolism, sterol metabolism, etc., and immune response, such as adaptive immune response, T cell activation, B cell activation, etc., (Fig. 4A, B, Fig. S4A, Table S6). These results suggested that the IVC condition may trigger elevated metabolism and immune response and resulted in compromised embryo quality. On the other hand, in IVV non-TE, genes related to development, transport, and cellular response were highly enriched (Fig. 4A, C, Fig. S4B, Table S6). It is noteworthy that we also found that several GO terms related to cell division, gene and protein regulations were highly enriched in IVC non-TE, whereas a number of apoptosis related GO terms were enriched in IVV non-TE (table S6).

**Fig. 4.**
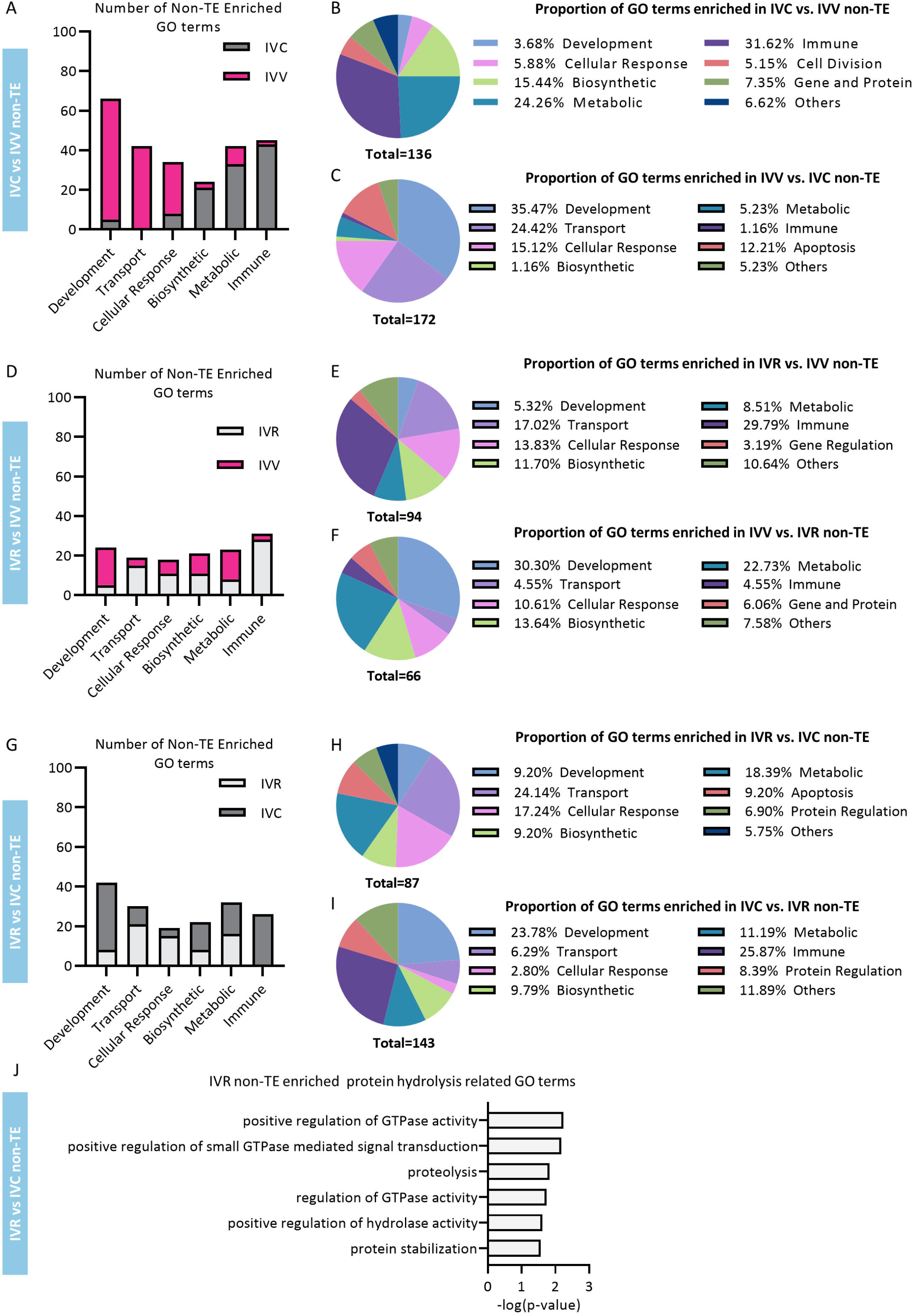
Differentially enriched GO terms in IVV, IVC, and IVR-derived non-TE lineage. A. Category and number of enriched GO terms in non-TE lineage when comparing IVC and IVV derived samples. B. Category and proportions of enriched GO terms in IVC-derived non-TE lineage when comparing to IVV derived samples. C. Category and proportions of enriched GO terms in IVV-derived non-TE lineage when comparing to IVC derived samples. D. Category and number of enriched GO terms in non-TE lineage when comparing IVR and IVV derived samples. E. Category and proportions of enriched GO terms in IVR- derived non-TE lineage when comparing to IVV derived samples. F. Category and proportions of enriched GO terms in IVV-derived non-TE lineage when comparing to IVR derived samples. G. Category and number of enriched GO terms in non-TE lineage when comparing IVR and IVC derived samples. H. Category and proportions of enriched GO terms in IVR-derived non-TE lineage when comparing to IVC derived samples. I. Category and proportions of enriched GO terms in IVC-derived non-TE lineage when comparing to IVR derived samples. J. Protein hydrolysis related GO terms enriched in IVR derived non-TE lineage when comparing to IVC derived samples.

When comparing IVR and IVV embryos, 1290 upregulated genes and 353 downregulated genes were found in the IVR-TE, while 346 genes were upregulated, and 337 genes were downregulated in the IVR-non-TE (Fig. S3C). GO analysis of non-TE cells revealed less differentially regulated GO terms between IVR and IVV (total = 160) when compared with the number of GO terms identified during IVC and IVV comparison (total = 308), suggesting IVR embryos may be more similar to IVV embryos (Fig 4D-F). However, IVR embryos still had large proportion of enriched GO terms related to immune response and very few GO terms related to development, suggesting the embryo quality is still compromised (Fig. 4D-F, Fig. S4C, D Table S8). Unlike IVC embryos, the number of GO terms related to biosynthetic process and cellular response were quite similar between IVR and IVV embryos (Fig. 4D-F), which indicated that IVR derived embryos were more resemble to embryos derived in vivo. Unsurprisingly, a high proportion of GO terms enriched in IVR was related to transmembrane transport due to the reduced nutrient level in medium, and metabolic process related GO terms were in fact more abundant in IVV embryos (Fig. 4D-F).

### IVR and IVC embryos have different proteolysis activity

The direct comparison of IVR and IVC non-TE revealed that IVC embryos had more genes enriched in GO terms associated with development and immune response, whereas, IVR embryos had more GO terms associated with transport and cellular response (Fig. 4G-I, Fig. S4E, F).The number of GO terms associated with metabolic process were similar between IVR and IVC embryos (Fig. 4G). An interesting observation is that both IVR and IVC embryos had a subset of GOs related to protein regulation (Table S7). The top GO terms enriched in IVR embryos in this category are positive regulation of GTPase activity, proteolysis, regulation of GTPase activity, and positive regulation of hydrolase activity (Fig. 4J). While the GO terms enriched in IVC embryos include negative regulation of peptidase activity, negative regulation of hydrolase activity, negative regulation of proteolysis, etc. (Fig. S4G). This result pointed to more active protein hydrolysis activity in IVR embryos, which is likely due to the conflict between the increased need of metabolic materials during lineage segregation and the insufficient nutrient level in environment.

### Analysis of transitional (C2) cells confirmed that IVR embryos are more similar to IVV embryos

As a peculiar lineage emerged during blastocyst stage, the transitional cluster (C2) was identified to have bipotential to either ICM or TE lineage, and most of the “ICM” cells stay in this status when cultured in vitro, which indicates that the transcriptomic features of these cells might determine the developmental potential of the embryos. As the reason for that, we sought to compare the DEGs among three treatments in this particular cell population. By merging the comparisons among all the gene sets, two large gene sets were identified, a total of 313 genes were commonly upregulated in IVV when compared to IVC and IVR groups (Figure 5A, orange bar), and a total of 296 genes were downregulated in IVC when compared to IVV and IVR groups (Fig. 5A, red bar), reflecting the difference between in vivo and in vitro culture conditions and also unveiling that IVR derived blastocysts were more similar to IVV derived blastocysts. The following GO/KEGG analysis showed both groups of genes were involved in mediating essential developmental processes and critical signaling pathways (Fig. 5B, table S9), for example, genes highly expressed in IVV group were enriched in regulating embryonic morphology, stem cell development, and apoptotic process, which were crucial for embryonic development and pluripotency of ICM cells. Furthermore, genes downregulated in IVC were involved in mediating several key pathways, including Notch, Ras, and Hippo signaling, which affecting cell polarity, embryonic cell differentiation, cell metabolism etc (Reichrath and Reichrath, 2020; Sonnen and Janda, 2021; Wu and Guan, 2021). These findings further implied that IVC and IVV embryos had the biggest difference, and IVR embryos were more similar to IVV embryos than IVC embryos.

**Fig. 5.**
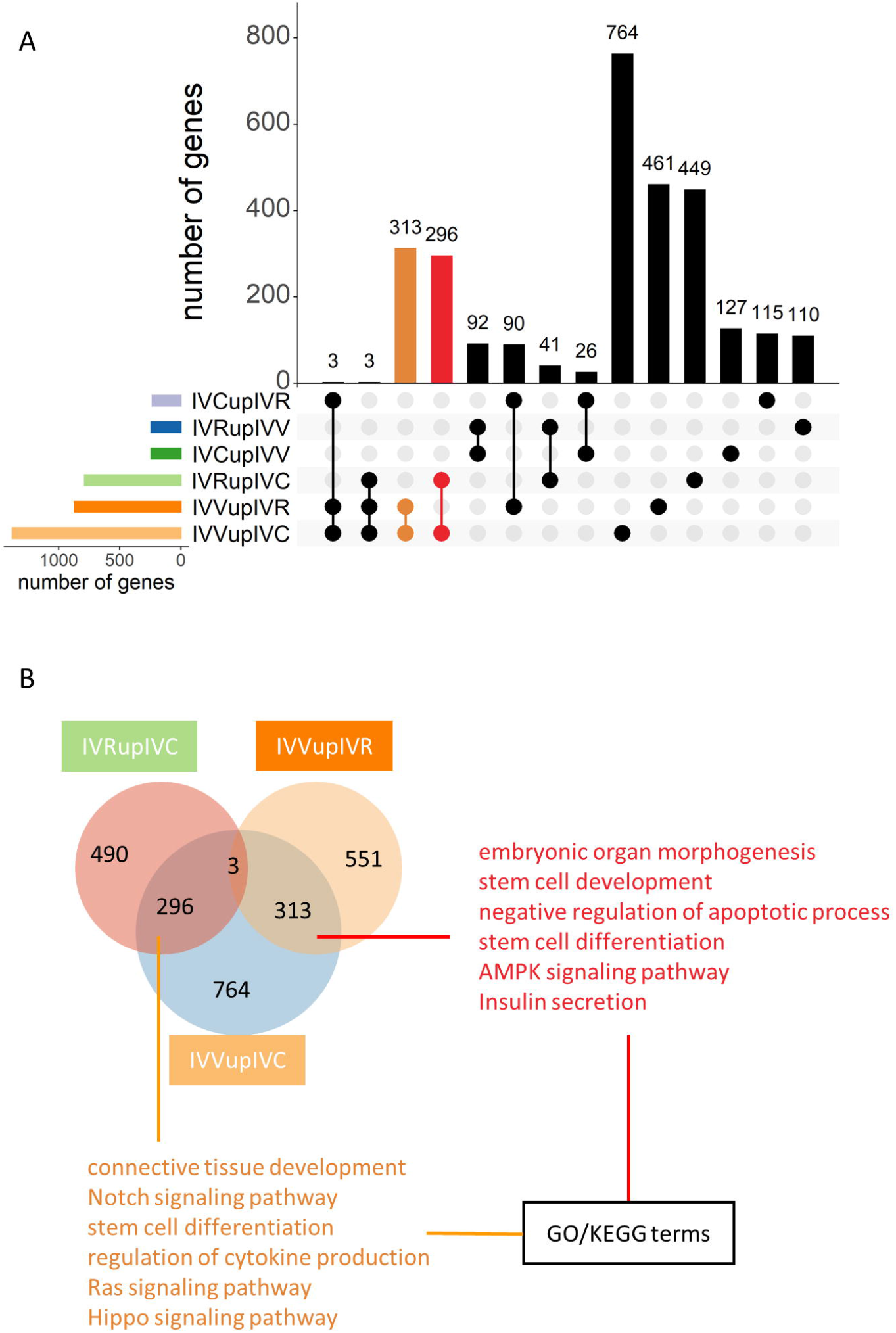
Comparison of transitional cell cluster (C2) derived from distinct environments. A. Upset graph showing the DEGs from samples in transitional stage (C2 cluster) between each two treatments. When compared to IVR, number of genes highly expressed in IVC was listed at first panel below the diagram and marked as IVCupIVR, so do other panels. The red bar showed genes downregulated in IVC when compared to both IVV and IVR. The orange bar showed genes highly expressed in IVV when compared to both IVC and IVR. B. Venn diagram showing the highly expressed genes in IVR when compared with IVC, as well as the highly expressed genes in IVV when compared with IVR and IVC, respectively. Specific GO/KEGG terms listed were enriched from two groups of genes as marked, respectively.

### Analysis of TE cells reveals additional information on how in vitro culture impact embryo quality

We also performed GO analysis on TE lineage, and unsurprisingly, the comparison between IVC and IVV embryos identified the largest number of enriched GO terms among all three comparisons (IVC vs. IVV, IVR vs. IVV, and IVR vs. IVC) (Fig.6A-H). A large number of differentially enriched GO terms between IVC and IVV TE were related to development, suggesting the TE differentiation under IVC condition is also compromised (Fig. 6A-C). Intriguingly, the metabolic process and immune response were more active in the IVV-TE, and transport was more active in IVC-TE (Fig. 6A-C, Fig. S5A, B), which are opposite to the observations we found in the non-TE. The differences between IVR and either IVV or IVC embryos were less distinctive (Fig. 6D, G). In consistent with what we found in the non-TE, many enriched GO terms in IVR-TE were associated with transport (Fig. 6D, E, G, H, Fig. S5C, E). We also found a group of GO terms related to homeostasis were downregulated in IVR-TE, when compared with IVV-TE, such as cellular monovalent inorganic cation homeostasis, cellular iron ion homeostasis, regulation of intracellular pH, etc (Fig. S5D). This could be explained as the side effect of the elevated transmembrane transportation. Another interesting finding is the negative regulation of cell growth was enriched in both IVR- and IVV-TE compared to the IVC-TE (Table S6, S7), suggesting that IVV and IVR embryos HAD controlled TE proliferation. This is in agreement with that embryos with lower ICM/TE ratio had compromised developmental potential (Qu et al., 2017).

**Fig. 6.**
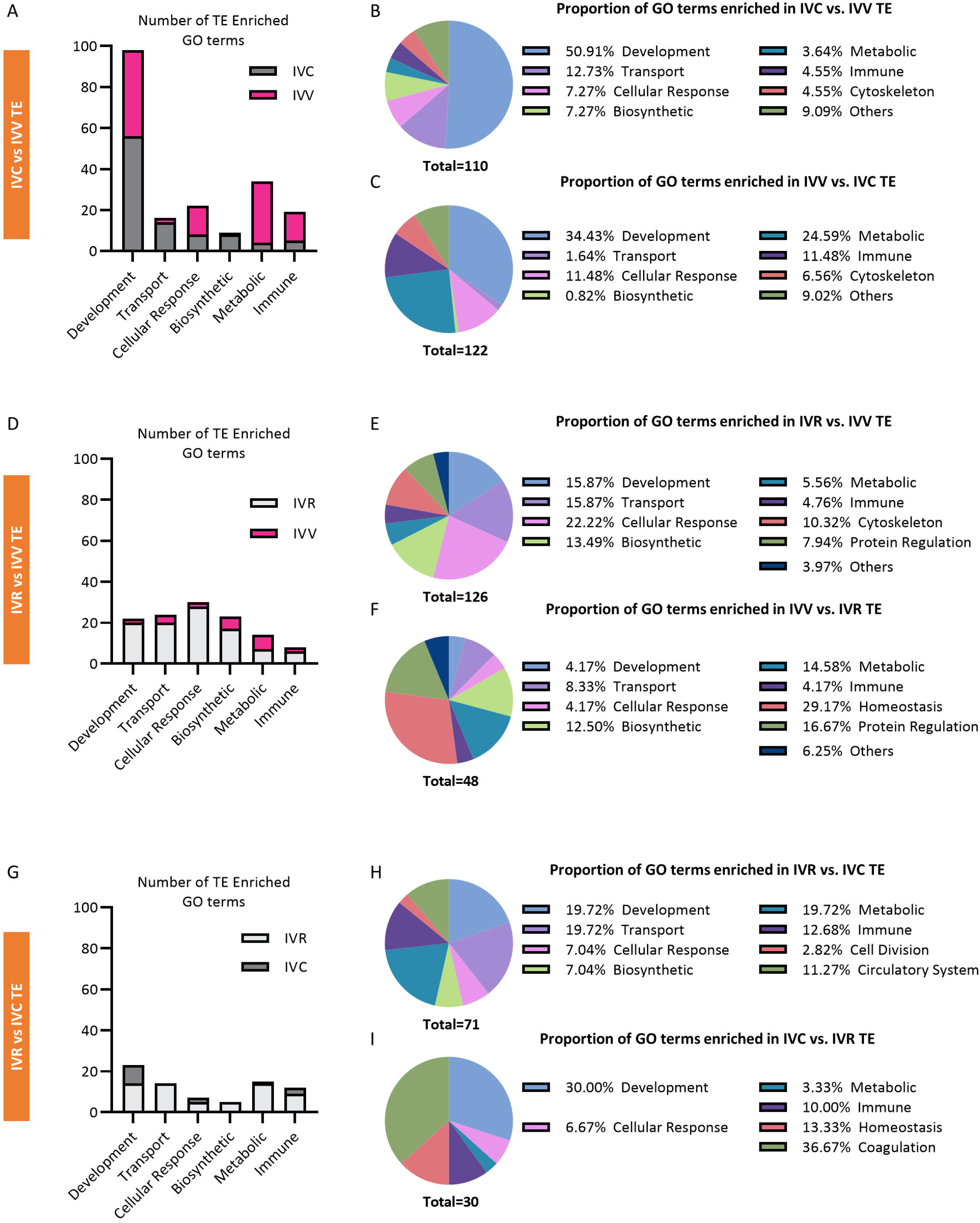
Differentially enriched GO terms in IVV, IVC, and IVR-derived TE lineage. A. Category and number of enriched GO terms in TE lineage when comparing IVC and IVV derived samples. B. Category and proportions of enriched GO terms in IVC-derived TE lineage when comparing to IVV derived samples. C. Category and proportions of enriched GO terms in IVV-derived TE lineage when comparing to IVC derived samples. D. Category and number of enriched GO terms in TE lineage when comparing IVR and IVV derived samples. E. Category and proportions of enriched GO terms in IVR-derived TE lineage when comparing to IVV derived samples. F. Category and proportions of enriched GO terms in IVV-derived TE lineage when comparing to IVR derived samples. G. Category and number of enriched GO terms in TE lineage when comparing IVR and IVC derived samples. H. Category and proportions of enriched GO terms in IVR-derived TE lineage when comparing to IVC derived samples. I. Category and proportions of enriched GO terms in IVC-derived TE lineage when comparing to IVR derived samples.

## Discussion

The segregation of TE and ICM is one of the most characteristic features during early embryonic development. ICM further differentiates into pluripotent EPI and the extraembryonic PE, which give rise to embryo proper and extra-embryonic cells (Rossant, 2018; Zhu and Zernicka-Goetz, 2020). The current understanding of this biological process is mostly based on studies from the mice model. Mice blastomeres become polarized at the morula stage. The outside polar cells generated following subsequent cell divisions form TE, while the inside apolar cells become enclosed by the polarized outer epithelium and form the ICM (Johnson and McConnell, 2004). This first lineage segregation is accompanied by the expression of key lineage markers, such as POU5F1 for ICM and CDX2 for TE, to establish each cell lineage (Nichols et al., 1998). The lineage segregation between EPI and PE in mouse blastocysts starts as early as E3.5, when the EPI specific marker Nanog, and PE specific marker Gata6 started to express and become mutually exclusive in the ICM in a random “salt and pepper” pattern (Chazaud et al., 2006). Although these two rounds of cell lineage segregation occur in every mammal, their spatiotemporal progressions are quite diverse. For example, the complete elimination of POU5F1 from the bovine TE cells requires several additional days after hatching (Berg et al., 2011). The segregation of the EPI and PE cell lineages was also different between mouse and bovine embryos, as Nanog knockout did not alter Gata6 expression and the initiation of PE differentiation in mouse embryos (Frankenberg et al., 2011), whereas knockout of NANOG in bovine embryos reduced GATA6 expression and impact the second lineage segregation (Ortega et al., 2020). Although GATA6-positive cells and NANOG-positive cells can be distinguished in day 8 bovine blastocysts (Kuijk et al., 2012), the lineage segregation of EPI and PE is probably much less restrictive at the blastocyst stage in bovine embryos compared to that in mouse. A recent study using the “on-gel” culture system to culture bovine embryos past blastocyst stage confirmed that the first and second lineage segregations were initiated after blastocyst cavitation and synchronously finished on day 9.5 embryos (Akizawa et al., 2021). Our scRNA-seq data reported here confirmed the lineage plasticity of the cells found in day 8 bovine blastocysts. The ICM cells (C3) have transcriptomic profiles featured with both EPI and PE markers identified during mouse development. We may be able to separate the precursor cells that tend to become either EPI or PE at this stage. However, it is quite clear that the segregation of EPI and PE has not readily occurred yet. The identification of the transitional cells (C2) with a mixed transcriptomic profile of ICM and TE even suggests the first lineage segregation of ICM and TE is still underway. The differentiation of the EPI, PE and TE may follow the “Waddington’s epigenetic landscape” (Wang et al., 2011), in which the undifferentiated ICM cells have the ability to transition to a more differentiated TE, however, the differentiation to TE may be irreversible. Therefore, it is important to separate the TE and non-TE and provide meaningful interpretation of how in vitro culture conditions affect the transcriptomic profile of each cell population in such a dynamic phase.

All the putative marker genes we examined here were universally expressed in all three clusters at different levels. As suggested by Akizawa et al. (Akizawa et al., 2021) and this study, the lineage specification of all three lineages is yet to accomplish in Day 8 bovine blastocysts, and many cells are in transitional state. Therefore, whether the differentially expressed genes in different cell clusters can be considered as lineage markers needs careful considerations. Nevertheless, the differentially expressed genes identified in our analysis did suggest the unique cell function of the precursor cells for each cell lineage we found in bovine blastocysts. This information may help us better define the cell identity and function of each cell type and proper culture conditions that encourage correct embryo development. In addition to the established lineage markers identified in other species, we discovered a new set of genes that may play important roles in lineages specification in bovine embryos. Immunoglobulin superfamily member 11 (IGSF11) was considered as a checkpoint in tumor immunotherapy, accompanied by multiple roles in regulating cell adhesion (Higashine et al., 2018), migration (Eom et al., 2012), and is essential for functional blood-testis barrier (Pelz et al., 2017). IGSF11 was preferentially expressed in ICM and might be involved in gap junction formation between cells. TDGF1 (Cripto-1) was found to express in both ICM and TE cells of mouse blastocyst and acts as mediator of implantation (Gershon et al., 2018). In addition, deficiency of TDGF1 leads to defects in mesoderm formation, resulting in heart defects (Barnes et al., 2016). Interestingly, in bovine blastocyst, TDGF1 was localized in ICM other than TE. Considering the existence of elongation stage in bovine, TDGF1 might emerge in TE much later. Cysteine dioxygenase type 1 (CDO1) is an integral enzyme catalyzing the metabolism of cysteine to sulfate. Its function was investigated in tumor (Kang et al., 2019) and human placenta (Dawson et al., 2020). We found that CDO1 was exclusively expressed in ICM, while its role in modulating physiological processes during early embryo development remains to be unveiled. LGALS3 encodes Galectin-3 which has been reported to have important effects on mouse embryonic implantation (Yang et al., 2012) and trophoblast development in human (Bozic et al., 2004). We further demonstrated its presence in bovine embryo as a validated TE marker. Another TE marker we validated was placenta-specific 8 (PLAC8). While it has already been used as a biomarker to evaluate bovine embryo quality (Ghanem et al., 2011; Machado et al., 2013; Silva et al., 2013), its distribution in blastocyst was not investigated. Here we showed that PLAC8 was preferentially expressed in TE and could serve as a specific lineage marker.

An important question this study wants to answer is how the in vitro condition influences the development of bovine blastocyst on transcriptomic level. The first thing to note is the impact of in vitro culture environment on cell lineage segregation. Regarding the non-TE, the most pluripotent ICM population (C3 cells) was almost exclusively derived from in vivo embryos, while most of the non-TE cells from IVC and IVR embryos were still in the transitional cell state (C2 cells). The IVV samples have much less transitional cells (C2 cells) when comparing to IVC or IVR samples, indicating the in vitro condition interrupted the cell segregation process, delaying the first cell fate determination, especially to ICM. In addition, GO term analysis comparing non-TE samples from IVV and IVC groups revealed that in IVC- derived non-TE, the more abundant genes were related to cellular metabolism and immune response in general, while IVV-derived non-TE cells are much more enriched in genes related to development, transport and cellular response. The “Quiet embryo” hypothesis suggested that preimplantation embryos are associated with a metabolism which is “quiet” rather than “active”, favoring embryo viability (Baumann et al., 2007). Our result of the non-TE lineage is in supportive of this theory that the more “active” in vitro produced embryos suggested a compromised development. This result suggested that, while the IVC blastocysts are still struggling in combating the adverse in vitro culture condition through increased metabolism and immune response, their IVV counterparts are well-positioned to prepare for the further lineage segregation and further development.

To examine how IVR condition influences the transcriptomic profile of bovine blastocysts, we first compared the IVR and IVV non-TE. In contrast to what we found in IVC embryos, we actually noticed less metabolic processes were enriched in IVR embryos compared to the IVV embryos. The downregulation of metabolism activity may directly connect to the upregulation of genes associated with cell survival and protein hydrolysis found in in IVR non-TE cells, indicating IVR condition may result in improved embryo quality through increased protein hydrolysis. Constant protein turnover is one of the major strategies utilized in maintaining cellular homeostasis (Toyama and Hetzer, 2013), and progressive loss of proteostasis is a hallmark of ageing that negatively impacts the proper cellular function and lifespan of individuals (Basisty et al., 2018). A large body of literature has confirmed that calorie restriction is, to date, the most effective non-genetic intervention to prevent age-associated diseases and extend longevity in most animal models (Lopez-Lluch and Navas, 2016). Interestingly, protein turnover rates are maintained at high levels in caloric restricted animals, and are implicated as one of the main mechanisms to limit oxidative damage and extend the lifespan of these animals (Tavernarakis and Driscoll, 2002). We believe similar mechanisms can apply here and explain why IVR embryos have higher viability and better developmental potential.

Another interesting finding from this comparison is that IVR embryos also had increased nutrient transport ability in both TE and non-TE. The increased nutrient transportation ability is probably necessary for embryos to acquire sufficient substrates in a reduced nutrient culture environment. This mirrors the findings that calorie restriction increases nutrient transport in animal and cell models (Casirola et al., 1997; Ferraris et al., 2001; Wang et al., 2017). Taken together, these data confirmed that the embryos cultured in IVR condition have better nutrient transport ability, have increased protein turnover to maintain better cellular homeostasis, therefore, reach to better cellular survival and function. The better quality of the IVR embryos was also confirmed by downregulation of the immune response related genes, suggesting that in the reduced nutrient condition, the cellular inflammatory status may be reduced.

Although we demonstrated the IVR condition may improve embryo quality, there were still significant transcriptomic differences between IVR and IVV embryos. In vitro environmental stress is not eliminated completely in the IVR condition, and the elevated immune response, suggesting increased inflammation, still exists in the IVR non-TE. During in vitro culture, embryos are subjected to variety of environmental stresses, including physical, chemical, oxidative, and energetic stresses, all of which can compromise embryo developmental potential (Summers and Biggers, 2003). Embryos respond to these environmental stresses with transcriptomic changes in metabolic and inflammatory signaling that may be necessary for embryo survival but affect their further development (Cagnone and Sirard, 2016). Another problem the IVR embryo experiencing is the over reactive transmembrane transport activities seen in both TE and non-TE. Few studies looked at nutrient transport capacity, its relationship with nutrient sensing, culture environment and how it impacts early embryo development. Future functional studies are needed to fill this knowledge gap. An interesting and surprising finding here is the downregulation of ion homeostasis related genes in the IVR embryos. The transport of substances across cell membranes may be the most fundamental activity of living cells. The adverse outcome of the overactive cellular transport is the difficulty to keep the appropriate cellular ion homeostasis and pH. In all, IVR embryos still need to keep up an increased nutrient transport activity and elevated immune response to battle the adverse in vitro culture condition, which take a heavy toll on embryo developmental potential.

In summary, we concluded the different features of blastocysts derived from three culture environments especially on non-TE and TE lineages. Notably, we found the major variation of embryo quality is from non-TE lineage. When using IVV as standard, IVC has increased metabolic, biosynthetic, and inflammatory activity, while development, cellular respond, and transmembrane transport ability are lower. IVR, however, has reduced metabolism, increased inflammation and transport activity, and importantly, it has comparable developmental potential as IVV (Fig. 7). As for TE part, compared to IVV, IVR and IVC have similar trends in metabolism, biosynthesis, and transportation, which indicates trophectoderm formation is barely affected by in vitro culture conditions.

**Fig. 7.**
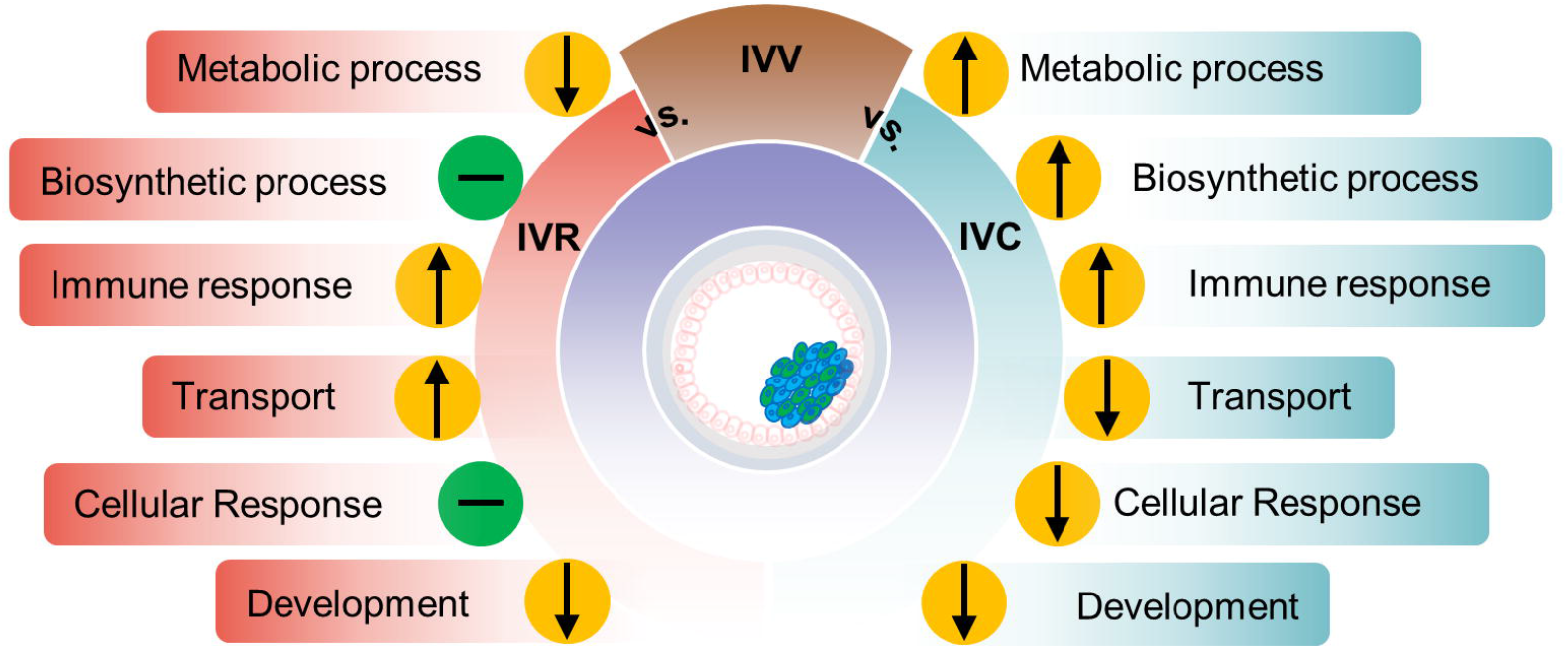
Proposed effects of in vitro culture environments on cellular functions in non-TE lineage. The conventional IVC condition resulted in increased embryo metabolic and biosynthetic processes and immune response, reduced transmembrane transport and reduced cellular response that contribute to the compromised development of IVC blastocysts. The IVR condition was able to maintain embryo biosynthetic process and cellular response activities similar to the in vivo counterparts. However, reduced metabolic process, increased immune response, and increased transmembrane transport activities suggest a different challenge the embryos have to face under the IVR condition.

For the first time, we described the transcriptomic profile of bovine blastocyst at the single cell level using scRNA-seq. This information is valuable in dissecting bovine lineage segregation at this critical development stage. The deep investigation of transcriptome by comparing samples from IVV, IVC and IVR derived blastocysts provided insights in how in vitro condition adversely impact embryo development and demonstrated the potential benefit of using the IVR condition to optimize bovine embryo IVP. This work paved a way for future functional studies to better understand how culture condition impact embryo development and further optimization of the in vitro culture condition.

## Materials and Methods

### Bovine in vitro and in vivo blastocyst production

For in vivo production, ovarian stimulation and embryo retrieval from cross bred cows were performed as previously described (Jiang et al., 2018; Ming et al., 2021). In brief, superovulation was achieved using five doses of intramuscular injections of FSH beginning 5 days after insertion of a Controlled Intravaginal Drug Release (CIDR) device. Two doses of prostaglandin F2 alpha were given along with the last two FSH treatments, followed by CIDR removal. Standing oestrus (Day 0) was seen approximately 48 h post-prostaglandin injection. GnRH was then administered at oestrus. Each cow was inseminated 12- and 24-h poststanding oestrus. Blastocysts were collected by routine non-surgical uterine flushing on days 8.

For in vitro production, ovaries were collected from a local slaughterhouse. Bovine cumulus-oocyte complexes (COCs) were aspirated from 2-6 mm antral follicles followed by 2-3 times wash in MOPS-buffered medium. The COCs were matured in a defined maturation medium containing 50 ng/ml recombinant murine EGF, 0.1 IU/ml recombinant human FSH (Merck & Co., Inc), 0.125 mg/ml recombinant human hyaluronan (Novozymes), and 2.5 mg/ml recombinant human albumin (AlbIX, Novozymes) for 22-24 hours. The matured COCs were washed in BO-IVF medium (IVF Bioscience) and transferred into fertilization drop containing BO-IVF medium (IVF Bioscience). Cryopreserved spermatozoa from a single bull were thawed and processed by pure sperm density gradient centrifugation (45%: 90%, Nidacon) followed 2 washes using MOPS-buffered medium. The purified spermatozoa were diluted with BO-IVF (IVF Bioscience) and added to fertilization drop containing matured COCs at the final concentration of 1 × 10^6^ spermatozoa/ml. After 18-20 hours of co-incubation, the presumptive zygotes were denuded by vortexing for 2.5 minutes to remove remaining cumulus cells and loosely bound spermatozoa. The denuded zygotes were randomly assigned to 20 µl culture drops of bovine Optimized Embryo Culture Medium 1(bOEC1) (Herrick et al., 2020). Embryo cleavage was assessed on day 3 (72h in bOEC1), and only embryos with more than 4-cells were washed and transferred to freshly prepared bOEC2 medium for additional 96 hour culture. Blastocysts were then evaluated under stereomicroscopy and only Grade 1 embryos by standards of the International Embryo Technology Society were selected for further study. Concentrations of salts (NaCl, KCl, KH2PO4, CaCl2-2H2O, MgSO4-7H2O, and 195 NaHCO3), antibiotics (gentamicin, 25 μg/ml), macromolecules (hyaluronan, 0.125 mg/mL), and growth factors (insulin, transferrin, and selenium, ITS) were kept the same to maintain consistent osmolarity and pH for both IVC and IVR medium. For IVR medium, nutrients (glucose/fructose, citrate, lactate, pyruvate, amino acids, vitamins, and EDTA) were diluted to 6.25% of the control IVC medium. The IVC medium was supplemented with 2.5 mg/mL lipid free BSA . The IVR medium was supplemented with 2.5 mg/ml lipid rich BSA (MP, 18052) with 5 mM L-carnitine supplementation.

### Isolation of single cells

Blastocysts were washed with DPBS containing 1 mg/ml polyvinylpyrrolidone (PBS-PVP) and transferred into 50 μl droplets of 0.1% protease (Qiagen) to remove the zona pellucida. Embryos were rinsed three times in PBS-PVP and confirmed to be free of contaminating cells. Prior to dissociation and single cell collection, half of the TE was removed to enrich the ICM cells. Single cells were isolated from embryos by incubating with TrypLE Express reagent for 10 min; and dissociated into single cells. We randomly picked about 10 cells per blastocyst, then transferred them separately into 0.2-mL PCR tubes (Eppendorf) containing cell lysis buffer and kept at −80 °C until library preparation.

### Single-cell RNA-seq library preparation and sequencing

The RNA-seq libraries were generated from individual cells by using the Smart-seq2 v4 kit with minor modification from manufacturer’s instructions. Briefly, individual cells were lysed, and mRNA was captured and amplified with the Smart-seq2 v4 kit (Clontech). After AMPure XP beads purification, the high-quality amplified RNAs were subject to library preparation using Nextera XT DNA Library Preparation Kit (Illumina) and multiplexed by Nextera XT Indexes (Illumina). The concentration of sequencing libraries was determined by using Qubit dsDNA HS Assay Kit (Life Technologies) and KAPA Library Quantification Kits (KAPA Biosystems). The size of sequencing libraries was determined by means of Agilent D5000 ScreenTape system at Tapestation 4200 system (Agilent). Pooled indexed libraries were then sequenced on the Illumina HiSeq X platform with 150-bp pair-end reads.

A total of 195 cells from 19 blastocysts were selected randomly to profile transcriptomes by RNA-seq using the Smart-seq2 protocol as described above. We generated approximately 22 million 150bp paired-end reads per sample.

### RNA-seq data processing

Multiplexed sequencing reads that passed filters were trimmed to remove low-quality reads and adaptors by TrimGalore (version 0.6.7) (-q 25 --length 20 --max_n 3 --stringency 3). QC for the data was performed on the FASTQ data using FastQC3 (v0.11.5). Trimmed reads were aligned to the bovine reference genome (ARS-UCD1.2) using HISAT25 (version 2.1.0) (--mp 6,2 --rdg 5,3--rfg 5,3). The reference genome was downloaded from ftp://ftp.ncbi.nlm.nih.gov/genomes/refseq/vertebrate_mammalian/Bos_taurus/latest_assembly_versions/GCF_002263795.1_ARS-UCD1.2/GCF_002263795.1_ARS-UCD1.2_genomic.fna.gz. The output SAM files were converted to BAM files and sorted using SAMtools6 (v1.3.1). Transcriptome assembly and gene expression quantification were performed using StringTie7 (version 1.3.3b) (-G Bos_taurus.ARS-UCD1.2.100.chr.gtf -A Sample_name.gene_ expression.txt). The genome annotation file was downloaded from ftp://ftp.ensembl.org/pub/release-100/gtf/bos_taurus/Bos_taurus.ARS-UCD1.2.100.gtf.gz. The read coverage (represented by read counts and TPM value) for each of the genes were generated by StringTie6 with the following parameters: -e -B - G GCF_002263795.1_ARS-UCD1.2_genomic.gtf -o Ballgown/Sample_ID/Sample_ID.gtf.

Clustering analysis was performed using R (version 3.3.3) package Monocle3. First, gene expression data were normalized by log and size factor to address depth differences. Linear dimensional reduction on the scaled data was performed using PCA. The top 10 principal components (PCs) were selected to represent a robust compression of the data. In order to visualize the cells in a two-dimensional space, UMAP was used to project the data into the X, Y plane (with parameter umap.n_neighbors = 15). Then, Louvain/Leiden community detection algorithm was applied to cluster all cells into different cell groups based on the UMAP result (configuration: k = 5, resolution = 0.001). Fine-tuning of the configuration was made in later part of the work and is noted in *Results*. PCA and tSNE were also used to visualize clustering results of all cells.

Differential gene expression analysis was performed by using R package edgeR with default parameters. The genes were deemed differentially expressed if they provided a false discovery rate of <0.01 and fold change >2. Volcano plot of DEGs was performed by using R package ggplot2. Biological pathways (denoted as Gene Ontology terms and KEGG pathways) over-represented among up- and down-regulated genes were identified using R package ClusterProfiler. P-value cutoff of 0.05 was used in selecting significant pathways. To further explore the function of DEGs in embryo and trophoblast development, GSEA was performed using R package GSEABase.

## Data availability

The raw FASTQ files are available at Gene Expression Omnibus (GEO) (https://www.ncbi.nlm.nih.gov/geo/) under the accession number GEO: GSE215409.

---

**Fig. S1.**
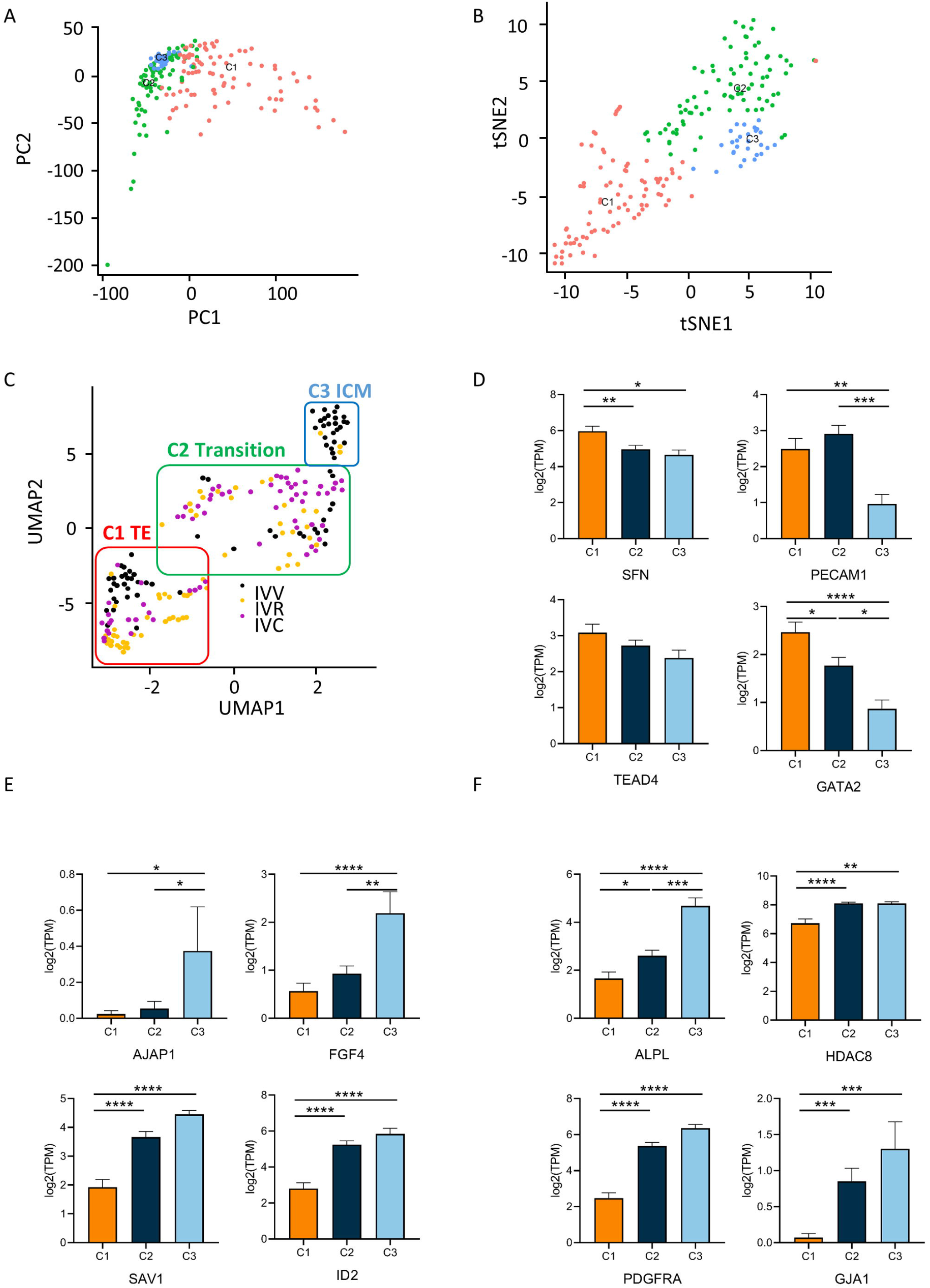
Cell cluster analysis with all samples. Distribution of samples in the two dimensional space, where the embeddings were computed by PCA (A) and t-SNE (B) using all expressed genes. C. Two-dimensional UMAP representation of cell clusters analyzed using all expressed genes, marked with corresponding lineage (TE, Transition, ICM) as well as culture conditions (IVV, IVR, IVC). Log2 TPM values of defined lineage marker genes in trophectoderm (D), EPI (E), primitive ectoderm (F). All data are presented as the mean ± SEM. * P < 0.05, ** P < 0.01, *** P < 0.001, **** P < 0.0001 from one-way ANOVA followed by Tukey’s multiple comparisons test.

**Fig. S2.**
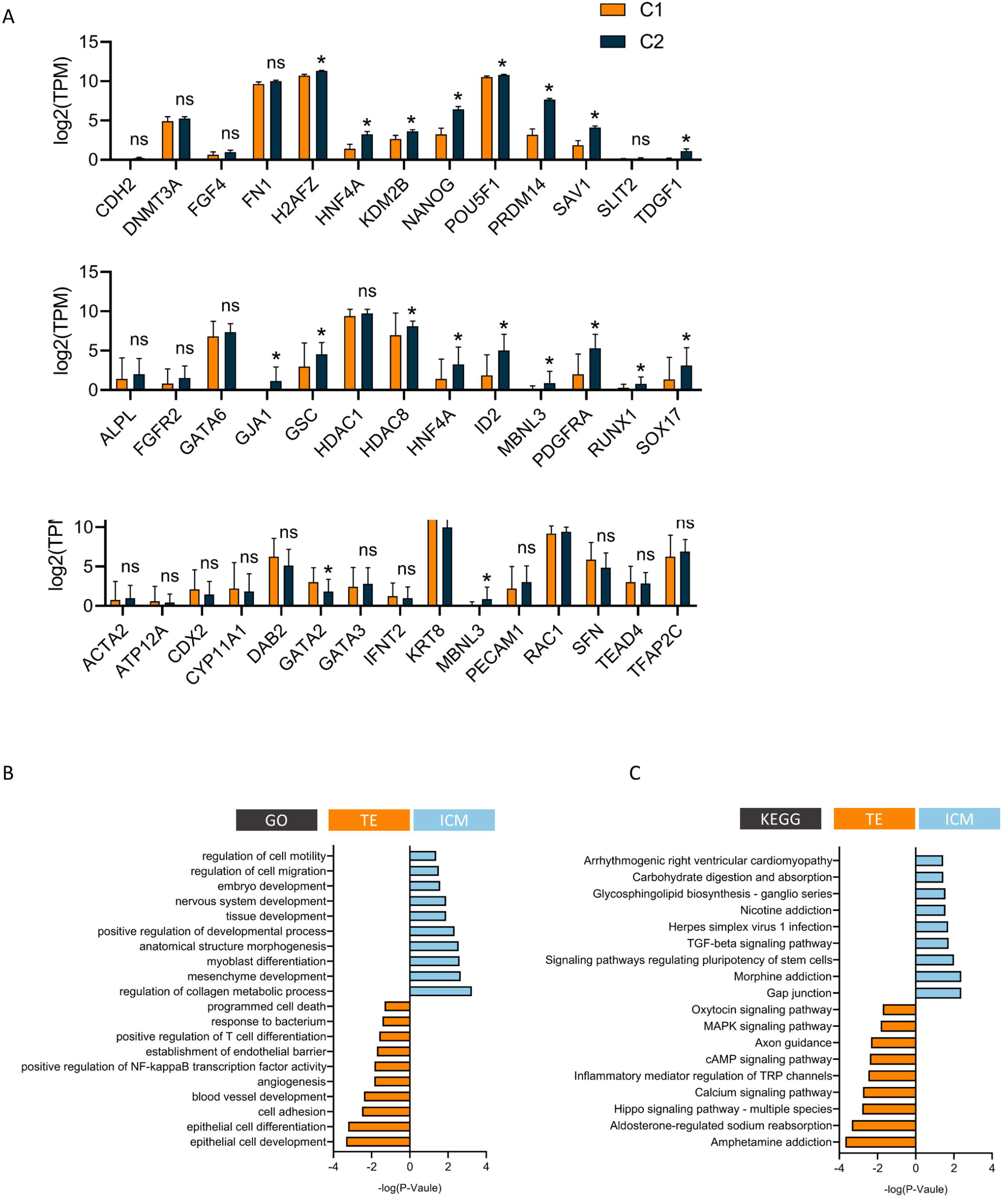
Differences between linages in in vivo derived bovine blastocyst. A. Log2 TPM values of defined lineage marker genes in C1 and C2 clusters derived from IVC embryo (top lane as EPI, middle lane as primitive ectoderm markers, bottom lane as markers trophectoderm markers). B. Represented gene ontology (GO) terms for ICM enriched genes (top) and TE enriched genes (bottom) in IVV embryos. C. Represented Kyoto Encyclopedia of Genes and Genomes (KEGG) terms for ICM enriched genes (top) and TE enriched genes (bottom) in IVV embryos. All data are presented as the mean ± SEM. * P < 0.05 from unpaired t-test.

**Fig. S3.**
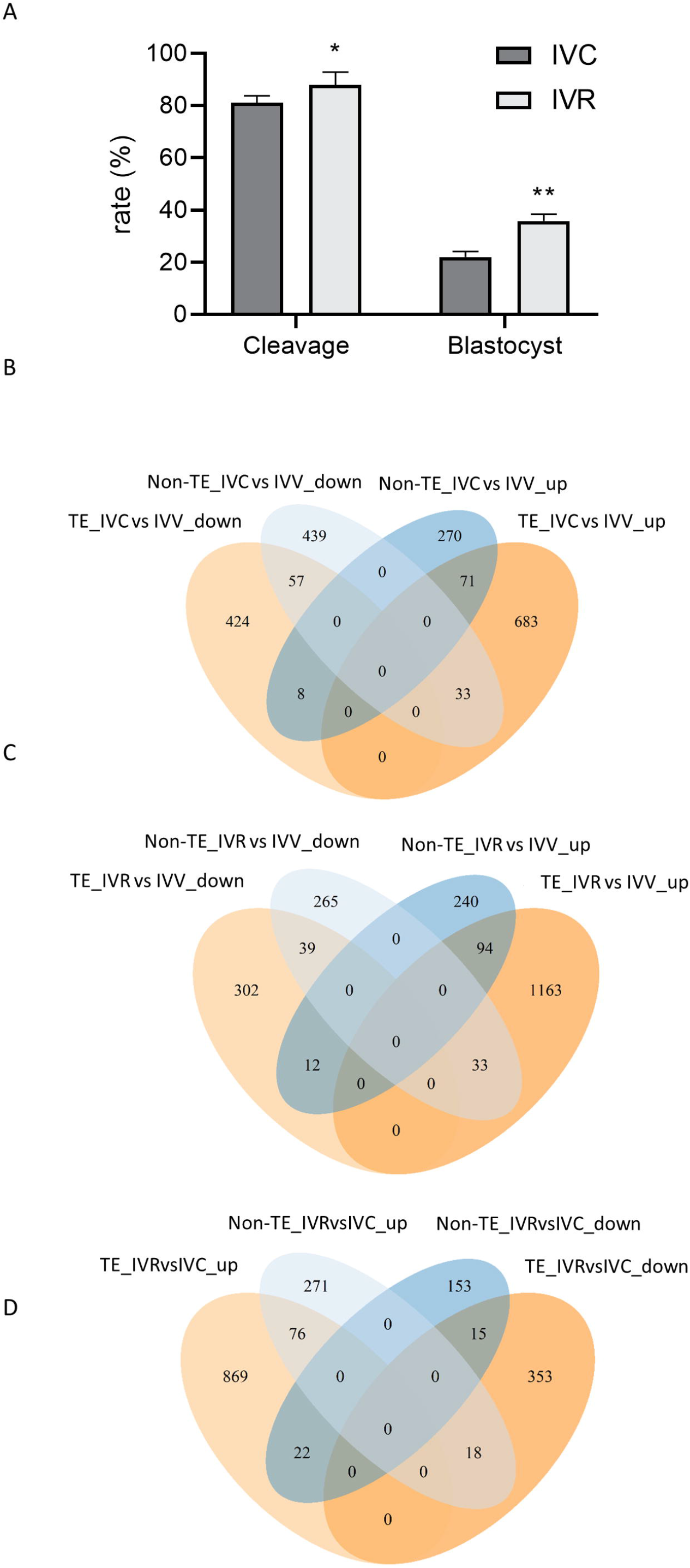
DEGs among separate lineages derived from different culture conditions. A. Embryo cleavage and blastocyst formation of bovine embryos culture in IVC (n=345 in total) and IVR (n=356 in total) conditions. This experiment was replicated five times. Embryo cleavage (p=0.02) and blastocyst formation (p=0.0002) were significantly between two IVC and IVR conditions. Data was analyzed as binomial proportions and compared using a Chi-squared test. B. Venn diagram of DEGs between IVV and IVC in both lineages. C. Venn diagram of DEGs between IVR and IVV in both lineages. D. Venn diagram of DEGs between IVR and IVC in both lineages.

**Fig. S4.**
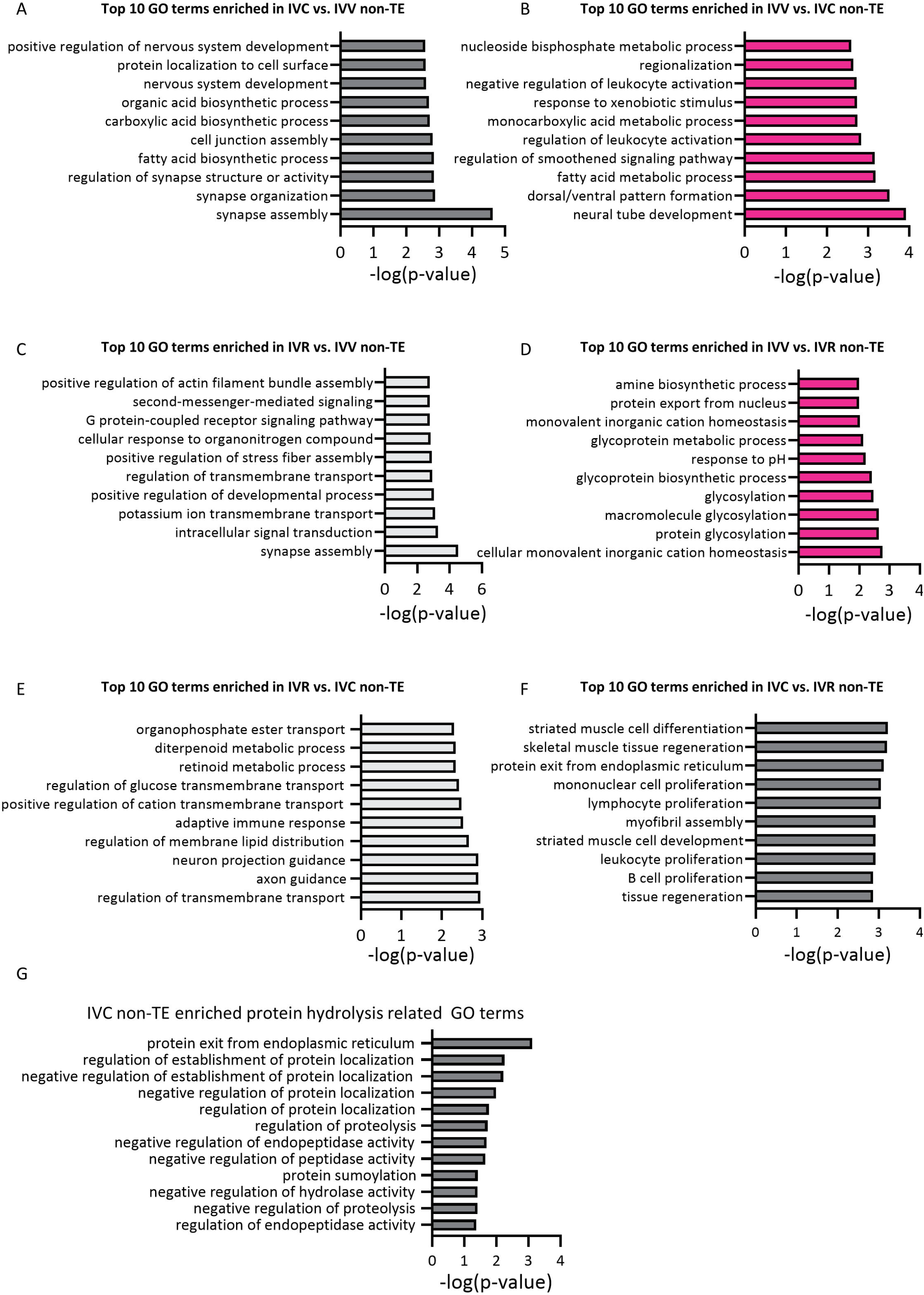
Top 10 GO terms enrichment in non-TE lineages. A. Top 10 GO terms enrichment in IVC non-TE lineage when comparing IVC and IVV derived samples. B. Top 10 GO terms enrichment in IVV non-TE lineage when comparing IVC and IVV derived samples. C. Top 10 GO terms enrichment in IVR non-TE lineage when comparing IVR and IVV derived samples. D. Top 10 GO terms enrichment in IVV non-TE lineage when comparing IVR and IVV derived samples. E. Top 10 GO terms enrichment in IVR non-TE lineage when comparing IVR and IVC derived samples. F. Top 10 GO terms enrichment in IVC non-TE lineage when comparing IVR and IVC derived samples. G. Protein hydrolysis related GO terms enriched in IVC derived non-TE lineage when comparing IVR and IVC derived samples.

**Fig. S5.**
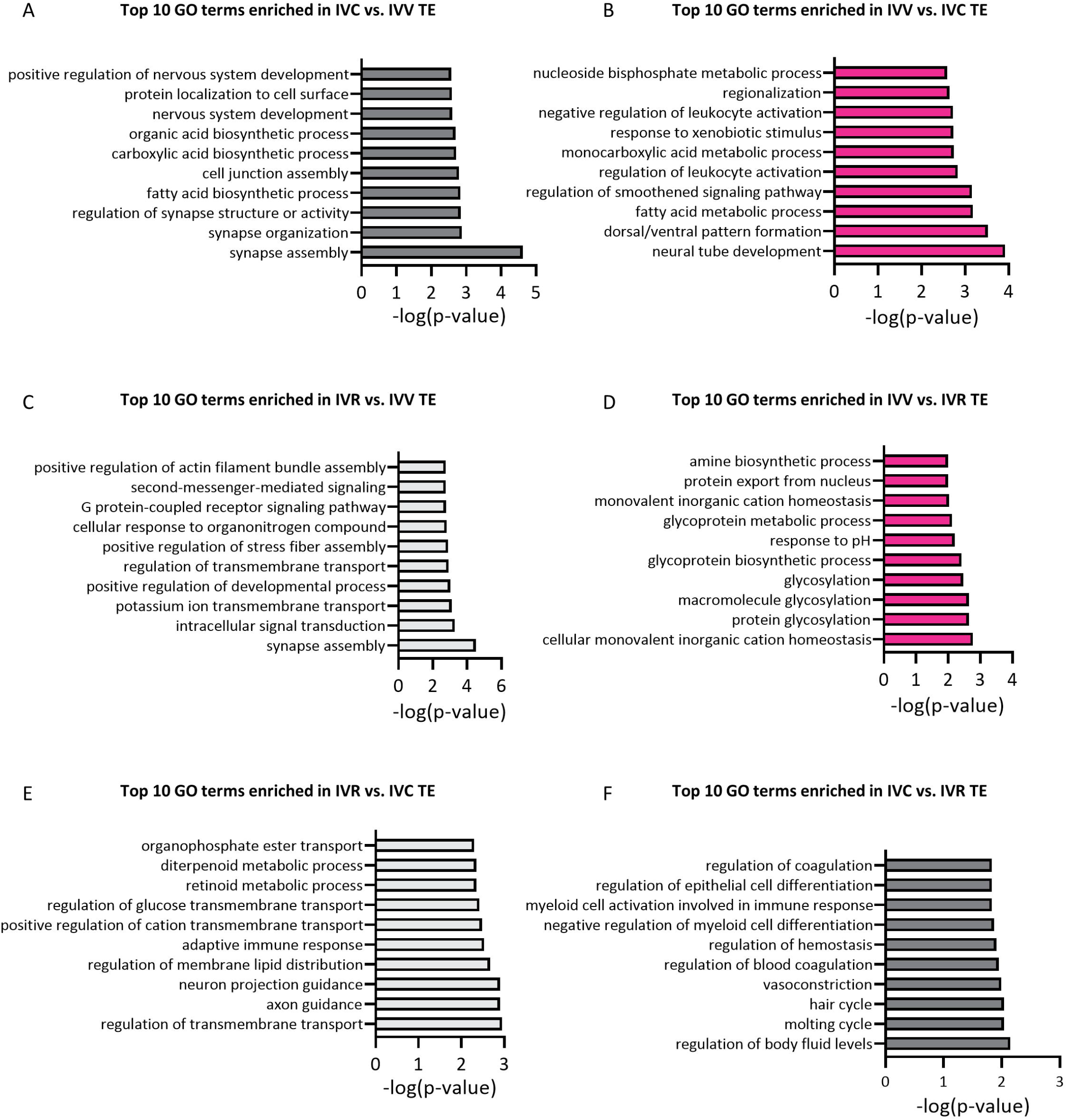
Top 10 GO terms enrichment in TE lineages. A. Top 10 GO terms enrichment in IVC TE lineage when comparing IVC and IVV derived samples. B. Top 10 GO terms enrichment in IVV TE lineage when comparing IVC and IVV derived samples. C. Top 10 GO terms enrichment in IVR TE lineage when comparing IVR and IVV derived samples. D. Top 10 GO terms enrichment in IVV TE lineage when comparing IVR and IVV derived samples. E. Top 10 GO terms enrichment in IVR TE lineage when comparing IVR and IVC derived samples. F. Top 10 GO terms enrichment in IVC TE lineage when comparing IVR and IVC derived samples.

## References

Akizawa, H., Saito, S., Kohri, N., Furukawa, E., Hayashi, Y., Bai, H., Nagano, M., Yanagawa, Y., Tsukahara, H., Takahashi, M., Kagawa, S., Kawahara-Miki, R., Kobayashi, H., Kono, T., Kawahara, M., 2021. Deciphering two rounds of cell lineage segregations during bovine preimplantation development. FASEB J 35, e21904.

Barnes, R.M., Harris, I.S., Jaehnig, E.J., Sauls, K., Sinha, T., Rojas, A., Schachterle, W., McCulley, D.J., Norris, R.A., Black, B.L., 2016. MEF2C regulates outflow tract alignment and transcriptional control of Tdgf1. Development 143, 774–779.

Basisty, N., Meyer, J.G., Schilling, B., 2018. Protein Turnover in Aging and Longevity. Proteomics 18, e1700108.

Baumann, C.G., Morris, D.G., Sreenan, J.M., Leese, H.J., 2007. The quiet embryo hypothesis: molecular characteristics favoring viability. Mol Reprod Dev 74, 1345–1353.

Berg, D.K., Smith, C.S., Pearton, D.J., Wells, D.N., Broadhurst, R., Donnison, M., Pfeffer, P.L., 2011. Trophectoderm lineage determination in cattle. Dev Cell 20, 244–255.

Bozic, M., Petronijevic, M., Milenkovic, S., Atanackovic, J., Lazic, J., Vicovac, L., 2004. Galectin-1 and galectin-3 in the trophoblast of the gestational trophoblastic disease. Placenta 25, 797–802.

Cagnone, G., Sirard, M.A., 2016. The embryonic stress response to in vitro culture: insight from genomic analysis. Reproduction 152, R247–R261.

Casirola, D.M., Lan, Y., Ferraris, R.P., 1997. Effects of changes in calorie intake on intestinal nutrient uptake and transporter mRNA levels in aged mice. J Gerontol A Biol Sci Med Sci 52, B300–310.

Chazaud, C., Yamanaka, Y., Pawson, T., Rossant, J., 2006. Early lineage segregation between epiblast and primitive endoderm in mouse blastocysts through the Grb2-MAPK pathway. Dev Cell 10, 615–624.

Dawson, P.A., Weerasekera, S.J., Atcheson, R.J., Twomey, S.A., Simmons, D.G., 2020. Molecular analysis of the human placental cysteine dioxygenase type 1 gene. Mol Genet Metab Rep 22, 100568.

Eom, D.S., Inoue, S., Patterson, L.B., Gordon, T.N., Slingwine, R., Kondo, S., Watanabe, M., Parichy, D.M., 2012. Melanophore migration and survival during zebrafish adult pigment stripe development require the immunoglobulin superfamily adhesion molecule Igsf11. PLoS Genet 8, e1002899.

Ermisch, A.F., Herrick, J.R., Pasquariello, R., Dyer, M.C., Lyons, S.M., Broeckling, C.D., Rajput, S.K., Schoolcraft, W.B., Krisher, R.L., 2020. A novel culture medium with reduced nutrient concentrations supports the development and viability of mouse embryos. Sci Rep 10, 9263.

Fan, X., Tang, D., Liao, Y., Li, P., Zhang, Y., Wang, M., Liang, F., Wang, X., Gao, Y., Wen, L., Wang, D., Wang, Y., Tang, F., 2020. Single-cell RNA-seq analysis of mouse preimplantation embryos by third-generation sequencing. PLoS Biol 18, e3001017.

Ferraris, R.P., Cao, Q.X., Prabhakaram, S., 2001. Chronic but not acute energy restriction increases intestinal nutrient transport in mice. J Nutr 131, 779–786.

Frankenberg, S., Gerbe, F., Bessonnard, S., Belville, C., Pouchin, P., Bardot, O., Chazaud, C., 2011. Primitive endoderm differentiates via a three-step mechanism involving Nanog and RTK signaling. Dev Cell 21, 1005–1013.

Gershon, E., Hadas, R., Elbaz, M., Booker, E., Muchnik, M., Kleinjan-Elazary, A., Karasenti, S., Genin, O., Cinnamon, Y., Gray, P.C., 2018. Identification of Trophectoderm-Derived Cripto as an Essential Mediator of Embryo Implantation. Endocrinology 159, 1793–1807.

Ghanem, N., Salilew-Wondim, D., Gad, A., Tesfaye, D., Phatsara, C., Tholen, E., Looft, C., Schellander, K., Hoelker, M., 2011. Bovine blastocysts with developmental competence to term share similar expression of developmentally important genes although derived from different culture environments. Reproduction 142, 551–564.

Herrick, J.R., Rajput, S., Pasquariello, R., Ermisch, A., Santiquet, N., Schoolcraft, W.B., Krisher, R.L., 2020. Developmental and molecular response of bovine embryos to reduced nutrients in vitro. Reprod Fertil 1, 51–65.

Higashine, K., Hashimoto, K., Tsujimoto, E., Oishi, Y., Hayashi, Y., Miyamoto, Y., 2018. Promotion of differentiation in developing mouse cerebellar granule cells by a cell adhesion molecule BT-IgSF. Neurosci Lett 686, 87–93.

Jiang, Z., Lin, J., Dong, H., Zheng, X., Marjani, S.L., Duan, J., Ouyang, Z., Chen, J., Tian, X.C., 2018. DNA methylomes of bovine gametes and in vivo produced preimplantation embryos. Biol Reprod 99, 949–959.

Johnson, M.H., McConnell, J.M., 2004. Lineage allocation and cell polarity during mouse embryogenesis. Semin Cell Dev Biol 15, 583–597.

Kang, Y.P., Torrente, L., Falzone, A., Elkins, C.M., Liu, M., Asara, J.M., Dibble, C.C., DeNicola, G.M., 2019. Cysteine dioxygenase 1 is a metabolic liability for non-small cell lung cancer. Elife 8.

Krisher, R.L., Heuberger, A.L., Paczkowski, M., Stevens, J., Pospisil, C., Prather, R.S., Sturmey, R.G., Herrick, J.R., Schoolcraft, W.B., 2015. Applying metabolomic analyses to the practice of embryology: physiology, development and assisted reproductive technology. Reprod Fertil Dev 27, 602–620.

Kuijk, E.W., van Tol, L.T., Van de Velde, H., Wubbolts, R., Welling, M., Geijsen, N., Roelen, B.A., 2012. The roles of FGF and MAP kinase signaling in the segregation of the epiblast and hypoblast cell lineages in bovine and human embryos. Development 139, 871–882.

Liu, D., Chen, Y., Ren, Y., Yuan, P., Wang, N., Liu, Q., Yang, C., Yan, Z., Yang, M., Wang, J., Lian, Y., Yan, J., Zhai, F., Nie, Y., Zhu, X., Chen, Y., Li, R., Chang, H.M., Leung, P.C.K., Qiao, J., Yan, L., 2022. Primary specification of blastocyst trophectoderm by scRNA-seq: New insights into embryo implantation. Sci Adv 8, eabj3725.

Lopez-Lluch, G., Navas, P., 2016. Calorie restriction as an intervention in ageing. J Physiol 594, 2043–2060.

Machado, G.M., Ferreira, A.R., Pivato, I., Fidelis, A., Spricigo, J.F., Paulini, F., Lucci, C.M., Franco, M.M., Dode, M.A., 2013. Post-hatching development of in vitro bovine embryos from day 7 to 14 in vivo versus in vitro. Mol Reprod Dev 80, 936–947.

Maddox-Hyttel, P., Alexopoulos, N.I., Vajta, G., Lewis, I., Rogers, P., Cann, L., Callesen, H., Tveden-Nyborg, P., Trounson, A., 2003. Immunohistochemical and ultrastructural characterization of the initial post-hatching development of bovine embryos. Reproduction 125, 607–623.

Ming, H., Sun, J., Pasquariello, R., Gatenby, L., Herrick, J.R., Yuan, Y., Pinto, C.R., Bondioli, K.R., Krisher, R.L., Jiang, Z., 2021. The landscape of accessible chromatin in bovine oocytes and early embryos. Epigenetics 16, 300–312.

Mole, M.A., Coorens, T.H.H., Shahbazi, M.N., Weberling, A., Weatherbee, B.A.T., Gantner, C.W., Sancho-Serra, C., Richardson, L., Drinkwater, A., Syed, N., Engley, S., Snell, P., Christie, L., Elder, K., Campbell, A., Fishel, S., Behjati, S., Vento-Tormo, R., Zernicka-Goetz, M., 2021. A single cell characterisation of human embryogenesis identifies pluripotency transitions and putative anterior hypoblast centre. Nat Commun 12, 3679.

Negron-Perez, V.M., Zhang, Y., Hansen, P.J., 2017. Single-cell gene expression of the bovine blastocyst. Reproduction 154, 627–644.

Nichols, J., Zevnik, B., Anastassiadis, K., Niwa, H., Klewe-Nebenius, D., Chambers, I., Scholer, H., Smith, A., 1998. Formation of pluripotent stem cells in the mammalian embryo depends on the POU transcription factor Oct4. Cell 95, 379–391.

Ortega, M.S., Kelleher, A.M., O’Neil, E., Benne, J., Cecil, R., Spencer, T.E., 2020. NANOG is required to form the epiblast and maintain pluripotency in the bovine embryo. Mol Reprod Dev 87, 152–160.

Pelz, L., Purfurst, B., Rathjen, F.G., 2017. The cell adhesion molecule BT-IgSF is essential for a functional blood-testis barrier and male fertility in mice. J Biol Chem 292, 21490–21503.

Qiu, H., Schlegel, V., 2018. Impact of nutrient overload on metabolic homeostasis. Nutr Rev 76, 693–707.

Qu, P., Qing, S., Liu, R., Qin, H., Wang, W., Qiao, F., Ge, H., Liu, J., Zhang, Y., Cui, W., Wang, Y., 2017. Effects of embryo-derived exosomes on the development of bovine cloned embryos. PLoS One 12, e0174535.

Reichrath, J., Reichrath, S., 2020. Notch Signaling and Embryonic Development: An Ancient Friend, Revisited. Adv Exp Med Biol 1218, 9–37.

Rossant, J., 2018. Genetic Control of Early Cell Lineages in the Mammalian Embryo. Annu Rev Genet 52, 185–201.

Santos, E.C.D., Fonseca Junior, A.M.D., Lima, C.B., Ispada, J., Silva, J., Milazzotto, M.P., 2021. Less is more: Reduced nutrient concentration during in vitro culture improves embryo production rates and morphophysiology of bovine embryos. Theriogenology 173, 37–47.

Silva, C.F., Sartorelli, E.S., Castilho, A.C., Satrapa, R.A., Puelker, R.Z., Razza, E.M., Ticianelli, J.S., Eduardo, H.P., Loureiro, B., Barros, C.M., 2013. Effects of heat stress on development, quality and survival of Bos indicus and Bos taurus embryos produced in vitro. Theriogenology 79, 351–357.

Sonnen, K.F., Janda, C.Y., 2021. Signalling dynamics in embryonic development. Biochem J 478, 4045–4070.

Summers, M.C., Biggers, J.D., 2003. Chemically defined media and the culture of mammalian preimplantation embryos: historical perspective and current issues. Hum Reprod Update 9, 557–582.

Tavernarakis, N., Driscoll, M., 2002. Caloric restriction and lifespan: a role for protein turnover? Mech Ageing Dev 123, 215–229.

Toyama, B.H., Hetzer, M.W., 2013. Protein homeostasis: live long, won’t prosper. Nat Rev Mol Cell Biol 14, 55–61.

Um, S.H., D’Alessio, D., Thomas, G., 2006. Nutrient overload, insulin resistance, and ribosomal protein S6 kinase 1, S6K1. Cell Metab 3, 393–402.

Wang, H., Arias, E.B., Yu, C.S., Verkerke, A.R.P., Cartee, G.D., 2017. Effects of Calorie Restriction and Fiber Type on Glucose Uptake and Abundance of Electron Transport Chain and Oxidative Phosphorylation Proteins in Single Fibers from Old Rats. J Gerontol A Biol Sci Med Sci 72, 1638–1646.

Wang, J., Zhang, K., Xu, L., Wang, E., 2011. Quantifying the Waddington landscape and biological paths for development and differentiation. Proc Natl Acad Sci U S A 108, 8257–8262.

Wei, Q., Zhong, L., Zhang, S., Mu, H., Xiang, J., Yue, L., Dai, Y., Han, J., 2017. Bovine lineage specification revealed by single-cell gene expression analysis from zygote to blastocyst. Biol Reprod 97, 5–17.

Wu, Z., Guan, K.L., 2021. Hippo Signaling in Embryogenesis and Development. Trends Biochem Sci 46, 51–63.

Yan, L., Yang, M., Guo, H., Yang, L., Wu, J., Li, R., Liu, P., Lian, Y., Zheng, X., Yan, J., Huang, J., Li, M., Wu, X., Wen, L., Lao, K., Li, R., Qiao, J., Tang, F., 2013. Single-cell RNA-Seq profiling of human preimplantation embryos and embryonic stem cells. Nat Struct Mol Biol 20, 1131–1139.

Yang, H., Lei, C., Zhang, W., 2012. Expression of galectin-3 in mouse endometrium and its effect during embryo implantation. Reprod Biomed Online 24, 116–122.

Zhao, L., Long, C., Zhao, G., Su, J., Ren, J., Sun, W., Wang, Z., Zhang, J., Liu, M., Hao, C., Li, H., Cao, G., Bao, S., Zuo, Y., Li, X., 2022. Reprogramming barriers in bovine cells nuclear transfer revealed by single-cell RNA-seq analysis. J Cell Mol Med 26, 4792–4804.

Zhu, M., Zernicka-Goetz, M., 2020. Principles of Self-Organization of the Mammalian Embryo. Cell 183, 1467–1478.

